# An adaptable analysis workflow for characterization of platelet spreading and morphology

**DOI:** 10.1101/2020.01.31.928168

**Authors:** Jeremy A. Pike, Victoria A. Simms, Christopher W. Smith, Neil V. Morgan, Abdullah O. Khan, Natalie S. Poulter, Iain B. Styles, Steven G. Thomas

## Abstract

The assessment of platelet spreading through light microscopy, and the subsequent quantification of parameters such as surface area and circularity, is a key assay for many platelet biologists. Here we present an analysis workflow which robustly segments individual platelets to facilitate the analysis of large numbers of cells while minimising user bias. Image segmentation is performed by interactive learning and touching platelets are separated with an efficient semi-automated protocol. We also use machine learning methods to robustly automate the classification of platelets into different subtypes. These adaptable and reproducible workflows are made freely available and are implemented using the open source software KNIME and ilastik.

## 1. Introduction

Testing platelet function in response to genetic mutations, gene knockouts and pharmacological agents is a valuable and widely used assay in platelet research^1–8^. In these studies the analysis of platelet spreading, either by the calculation of adhesion levels, spread surface areas or morphological categorisation, is used as a measure of platelet function. As platelets are small cells typically two to four microns in diameter, a single light microscopy field of view (FOV) can capture 50 - 400 platelets. As such it is easy to acquire data for large populations of cells allowing for the identification of subtle changes. In addition, immunofluorescence based labelling allows quantitative measures of platelet area and morphology to be combined with analysis of protein sub-cellular localisation and organisation.

Despite this, the analysis of platelet spreading can be a laborious process, especially in large scale experiments, where many thousands of platelets over a range of conditions might need to be analysed. A common way of measuring platelet spread area is to manually draw around the outline of the cell^9^. However, this is an extremely slow process which limits its application to larger datasets.

A more efficient and typically less biased way to perform the analysis is to design an automated image analysis workflow (not machine learning based) which is automated using reproducible and preferably open-source software such as ImageJ/Fiji^10^. Such workflows typically employ simple filtering operations and thresholds on image intensity^7^. The free parameters of the workflow are then set ad-hoc and rarely perform well across large datasets. Moreover, these workflows are usually only applicable to images captured on a particular microscope, with cells stained, or imaged, under very specific conditions. The categorisation of platelets into sub-types based on spread morphology is typically performed manually, and is therefore time-consuming and highly susceptible to user bias. We present a simple, adaptable workflow which uses machine learning based techniques to overcome many of these limitations, and thus allows for the robust quantitative analysis of platelet spreading across different imaging modalities and laboratories.

## 2. Method

### 2.1 Workflow description

An overview of the workflow is presented in Figure 1. The first step is the segmentation of platelets from the background to produce binary (black and white) images. To do this we use a pixel classifier trained within the open-source software ilastik^11,12^. Briefly, various pixel-level features including smoothed intensity and edge indicators are measured and used to train a random forest classifier with two outcomes; signal and background. Training images should be selected across replicates and treatments to ensure the full variability within the dataset is captured. Having trained the pixel classifier within ilastik, it is run on the full dataset along with all subsequent analysis steps within KNIME^13^, another open-source data analysis platform.

**Figure 1.**
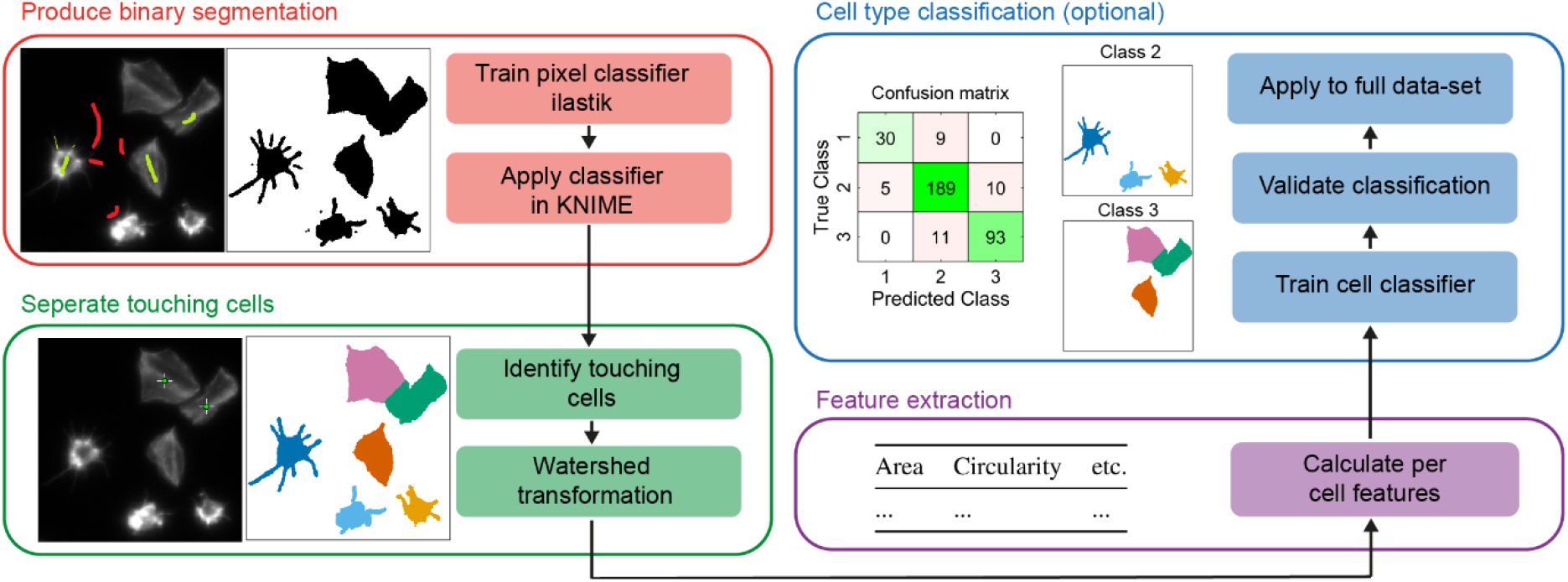
Overview of the proposed workflow for analysis of platelet spreading. First a pixel classifier is used to produce a binary segmentation mask. Next touching cells are manually annotated by clicking on their centre within KNIME and a watershed transform is used to establish the cell-cell boundaries. Per cell features are then calculated which can optionally be used to train a cell classifier. Validation of the classifier is achieved by reserving a proportion of the training data and visualised through a confusion matrix.

The second step is the separation of touching platelets. For this we chose to use a semi-automated approach where the researcher is asked to click on the centre of all touching platelets. These points are then used as seeds for a watershed transform which fills the binary images produced by the pixel classifier. This produces labelled segmentation images where each cell has a unique pixel value which facilitates the separation of touching cells. A connected component analysis is then used to calculate per platelet morphological features including area and circularity.

Finally, the population can be further interrogated by defining platelet morphological subtypes, for example unspread, partially spread and fully spread, and then using a machine learning approach to classify individual cells (objects). Again a random forest classifier is used, but it is trained using platelet morphological features, including area and circularity, as opposed to pixel level features like intensity. This quantifies the number of cells in each class and allows for the detailed morphological analysis of cells within a specific class. The corresponding workflows, and a detailed user guide can be found in the supplementary materials.

### 2.2 Testing and validation

To test and validate the workflow we chose to fluorescently label F-actin and image using wide-field microscopy, a commonly used approach. The cells were spread on either collagen or fibrinogen and treated either with dasatinib or a DMSO control (Figure 2). Dasatinib is a Src family kinase inhibitor and is known to reduce the efficiency of platelet spreading^14,15^. We chose to use a treatment with well-established effects as this provides a better benchmark for workflow validation. Three technical replicates with three fields of view per replicate were acquired. Further experimental details can be found in the supplementary materials.

**Figure 2.**
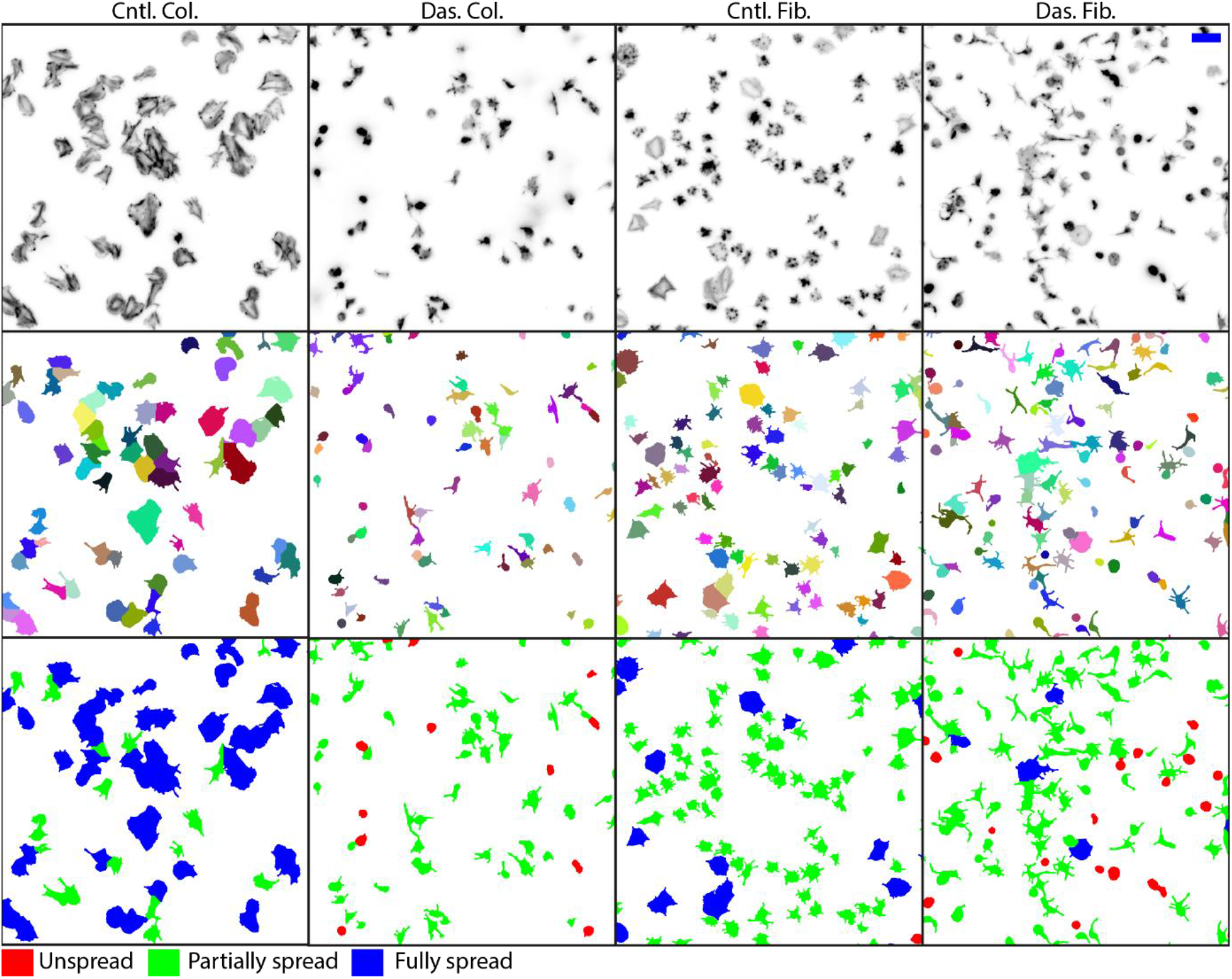
Representative cropped images and results from platelets seeded on either collagen (Col.) or fibrinogen (Fib.) and treated with either dasatinib (Das.) or a DMSO control (Cntl.). Top row shows a maximal projection of the raw data (inverted grey-scale look-up-table). This is used as the input for the analysis workflow. Middle row shows the individual platelet segmentations where each cell is a distinct colour. Bottom row shows the results of the object classifier where individual platelets are classified as either unspread (red), partially spread (green) or fully spread (blue). Scale bar 10 μm.

The images were acquired as z-stacks so these were pre-processed by finding the most in-focus image of the stack and then taking the maximal projection across this slice and the two slices either side (5 slices in total). This was done to limit out-of-focus contributions to the projections and improve subsequent segmentation. Vollath’s F4 measure was used as the focus metric,^16,17^ which is implemented using ImgLib2^18^, within the KNIME workflows provided. The 2D projections were then processed with the remaining workflow steps including classification into the following pre-defined categories; unspread, partially spread and fully spread. In total across all conditions 9655 platelets were segmented and analysed. Eight cropped images selected across replicates and conditions were used to train both the pixel and object classifiers.

To evaluate the performance of the object classifier twenty percent of the annotated platelets were reserved for validation. When using all measured morphological features the overall classification accuracy was found to be 90% and the corresponding confusion matrix is shown in Figure 3a. Classification accuracies of 71% and 77% were found when area or circularity were used as the only input. 87% accuracy was obtained using both area and circularity. This indicates that thresholds on area and circularity alone are not optimal for robust and accurate classification of platelets into sub-categories, and highlights the advantage of a machine learning approach which uses a larger number of different features.

**Figure 3.**
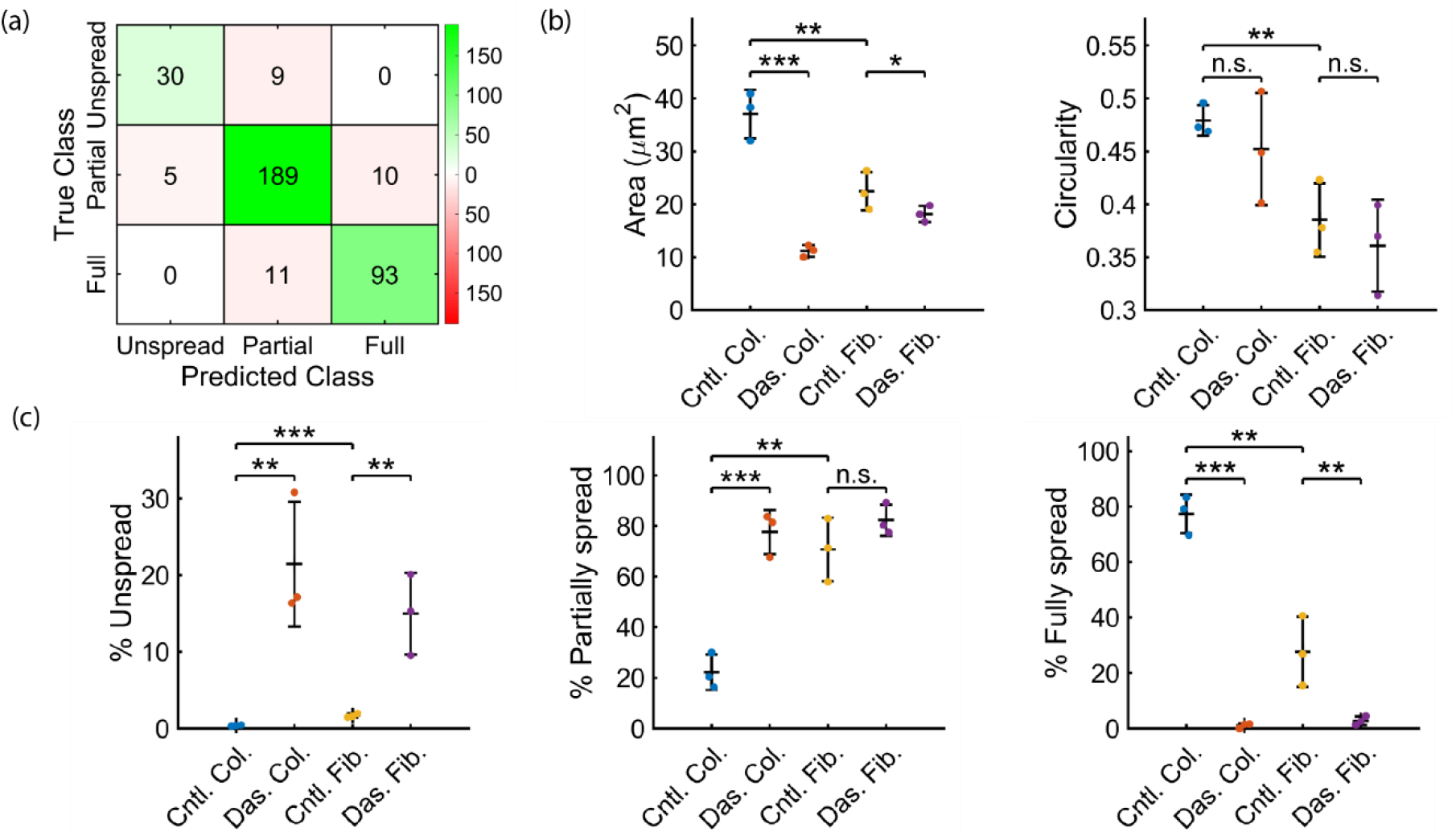
Summarised quantitative outputs of the analysis workflow. Platelets were seeded on either collagen (Col.) or fibrinogen (Fib.) and treated with either dasatinib (Das.) or a DMSO control (Cntl.) **(a)** A confusion matrix allows for visual evaluation of the object classifier. A proportion of the training data is reserved (here 20%) and the class predicted by the classifier is compared to the true class as defined by the manual annotation. On-diagonal classifications (green) represent agreement between the classifier and manual annotation, off-diagonal classifications (red) represent disagreement. **(b)** Mean platelet area and circularity calculated across all platelets in a replicate (N=3, mean 805 platelets per replicate). **(c)** Percentage of cells in each category; unspread, partially spread and spread. All statistical analyses by one-way Anova and subsequent pair-wise comparison by two-sample t-test with Bonforonni correction. ***P<0.001, **P<0.01, *P<0.05, error bars are mean ± s.d.

Figure 3b shows the measurements for mean platelet area and circularity. As expected there are significant differences in platelet area between the collagen and fibrinogen controls, and also with the dasatinib treatment on both substrates. For circularity the only significant difference observed is between the collagen and fibrinogen controls. Figure 3c shows the relative proportion of each class type. There are clear differences between conditions, highlighting the advantage of the classification approach in further delineating the platelet spread phenotype. For example, treatment with dasatinib dramatically reduces the percentage of fully spread platelets on both substrates.

## 3. Discussion

In this manuscript, we have described a semi-automated analysis workflow for the quantification of platelet spreading. We demonstrate that, following the training of a pixel classifier on a small subset of data, this method is able to accurately segment and quantify the spread surface area and circularity of platelets treated with dasatinib (which at the concentration used here blocks both Btk and Src family kinases^14^) on both collagen and fibrinogen surfaces. The workflow was able to appropriately identify and segment both isolated cells and platelets touching other platelets. Subtleties of platelet morphology such as filopodia were clearly identified and quantification of the parameters such as spread area were able to give a simple, robust overview of the effect of the inhibitor treatment on platelet spreading. Furthermore, an object classifier was used to group platelets into classes which allowed for deeper interrogation of the data.

A key advantage of the reported workflow is the ability to efficiently analyse large numbers of platelets (we routinely measure and classify 10,000+ platelets using the workflow) which allows robust statistical analyses to be performed. The power of this approach was demonstrated in a recent study of 55 samples from patients with bleeding of unknown cause^19^. We have used the workflow for both human and mouse platelets and it is applicable to a wide range of treatments (e.g. patient samples, gene knockouts, inhibitor studies, etc.). Therefore, this work presents a viable way to perform quick and accurate large scale analysis of spreading as a measure of platelet function while also minimising user bias.

When designing the workflow care was taken to ensure each step was robust to different imaging systems and sample preparations so as to be widely applicable. Provided it is re-trained, the ilastik pixel classifier will perform well across a range of stains and non-florescent imaging modalities, for example phase-contrast microscopy (Supplementary Figure 1). However, where feasible we recommend florescent staining to enhance the contrast between cells and background, and note that as with all image analysis processes, poor quality input data may result in incorrect classification or failure of the analysis. For object classification the classes are defined by the researcher and so can be changed to suit the biological question. Although it is important to note that classification will be more successful with classes that have a clearly distinct morphology.

With regards to this final point, it is important to check the classification performance on a subset of the annotated data reserved for this purpose. Performance can then be assessed though overall classification accuracy and confusion matrices (Figure 3a). Tools and instructions for this performance evaluation are including in the provided workflows and guidelines. Manually selected thresholds on parameters such as area and circularity were avoided as they are non-optimal and rarely robust across replicates and conditions. For both pixel and object classification, the training can be performed quickly with a small amount of data (typically 5 – 10 cropped images) to produce high quality results. Furthermore the graphical programming interface offered by KNIME means that researchers with no, or limited, programming experience can adapt these protocols for their specific needs.

The workflow is fully automated apart from the manual selection of touching platelets which is performed by the researcher within KNIME. Automated separation for other densely packed cell types is typically achieved by first segmenting nuclei which are then used as seeds to isolate the cytoplasm. This approach is not applicable to platelets which have no nuclei, hence the need to manually identify touching cells. Moreover platelet morphology can vary dramatically dependent on the surface coating and treatment which further complicates the task of automated separation. Further research will investigate if with sufficient training data, deep learning based methods can be used to robustly segment clustered platelets.

## 4. Conclusion

We present a semi-automated workflow that can be applied to segment, classify and analyse spread platelets. The workflow is adaptable and applicable to input images from a range of imaging modalities. Once trained the workflow can perform efficient analysis of large data sets and provides an unbiased measure of platelet spreading. These factors, along with the use of open source software, should allow for wide uptake by platelet researchers, who will be able to use these tools to perform robust analyses on large scale image data.

## Acknowledgements

The authors gratefully acknowledge Steve P Watson and the members of the Birmingham platelet group who provided many useful comments and advice. The work was funded by the Centre of Membrane Proteins and Receptors (COMPARE), Universities of Birmingham and Nottingham, Midlands, UK and from the British Heart Foundation through the Chair award (CH0/03/003) to Steve P Watson and a project grant (PG/15/114/31945) to Steven G Thomas.

## Supplementary materials and methods

### Platelet preparation and treatment

Platelets were isolated from human blood samples, which were donated by healthy volunteers. Human venous blood was drawn by venipuncture into sodium citrate and acid-citrate-dextrose solution. Whole blood was centrifuged at 200 × g for 20 minutes to obtain platelet rich plasma. 0.1 μg/ml prostacyclin was added to the platelet rich plasma and platelets were collected after centrifugation at 1000 × g for 10 minutes. The platelet pellet was re-suspended in acid-citrate-dextrose and modified Tyrode’s buffer containing 129 mM NaCl, 0.34 mM Na2HPO4, 2.9 mM KCl, 12 mM NaHCO3, 20 mM HEPES, 5 mM glucose, 1 mM MgCl2, pH 7.3 and 0.1 μg/ml prostacyclin was added to the washed platelets and centrifuged at 1000 × g for 10 minutes to be washed. The platelet pellet was re-suspended in modified Tyrode’s buffer to a concentration of 2 × 10^8^ platelets/ml and left to rest for at least 30 minutes before further dilution to 2 × 10^7^ platelets/ml prior to being used in spreading experiments.

Coverslips were coated with 100 μg/ml fibrinogen or 10 μg/ml collagen and left overnight at 4 °C. The unbound fibrinogen and collagen were removed and the coverslips were blocked using 5 mg/ml bovine serum albumin (BSA) in PBS for 1 hour and washed with PBS prior to use. All spreading experiments were performed in the presence of 2 U/ml apyrase and 10 μM indomethacin. 2 × 10^7^ platelets/ml washed platelets were either incubated for 10 minutes at 37 °C in the presence of 10 μM dasatinib or DMSO, prior to 45 minutes of spreading on pre-coated coverslips at 37 °C.

### Imaging

Platelets were fixed after spreading using 10% neutral buffered formalin solution for 10 minutes at room temperature. 0.1% Triton X-100 was added for 5 minutes at room temperature prior to platelets being washed with PBS and incubated with Alexa488-phalloidin for 1 hour at room temperature to stain the filamentous (F)-actin fibres present within the platelets for the observation of platelet morphology. The actin stained platelets were mounted onto glass slides using hydromount. Images were acquired using an Axio Observer 7 inverted epifluorescence microscope (Carl Zeiss Microscopy) with Definite Focus 2 autofocus, 63x 1.4 NA oil immersion objective lens, Colibri 7 LED illumination source, Hammamatsu Orca Flash 4 V2 sCMOS camera, Filter set 38 for Alexa488 and DIC optics. LED power and exposure time were chosen as appropriate for each set of samples but kept the same within each experiment. Using Zen 2.3 Pro software. Three image stacks (step size 0.3 μm) were taken per coverslip.

---

**Supplementary Figure 1.**
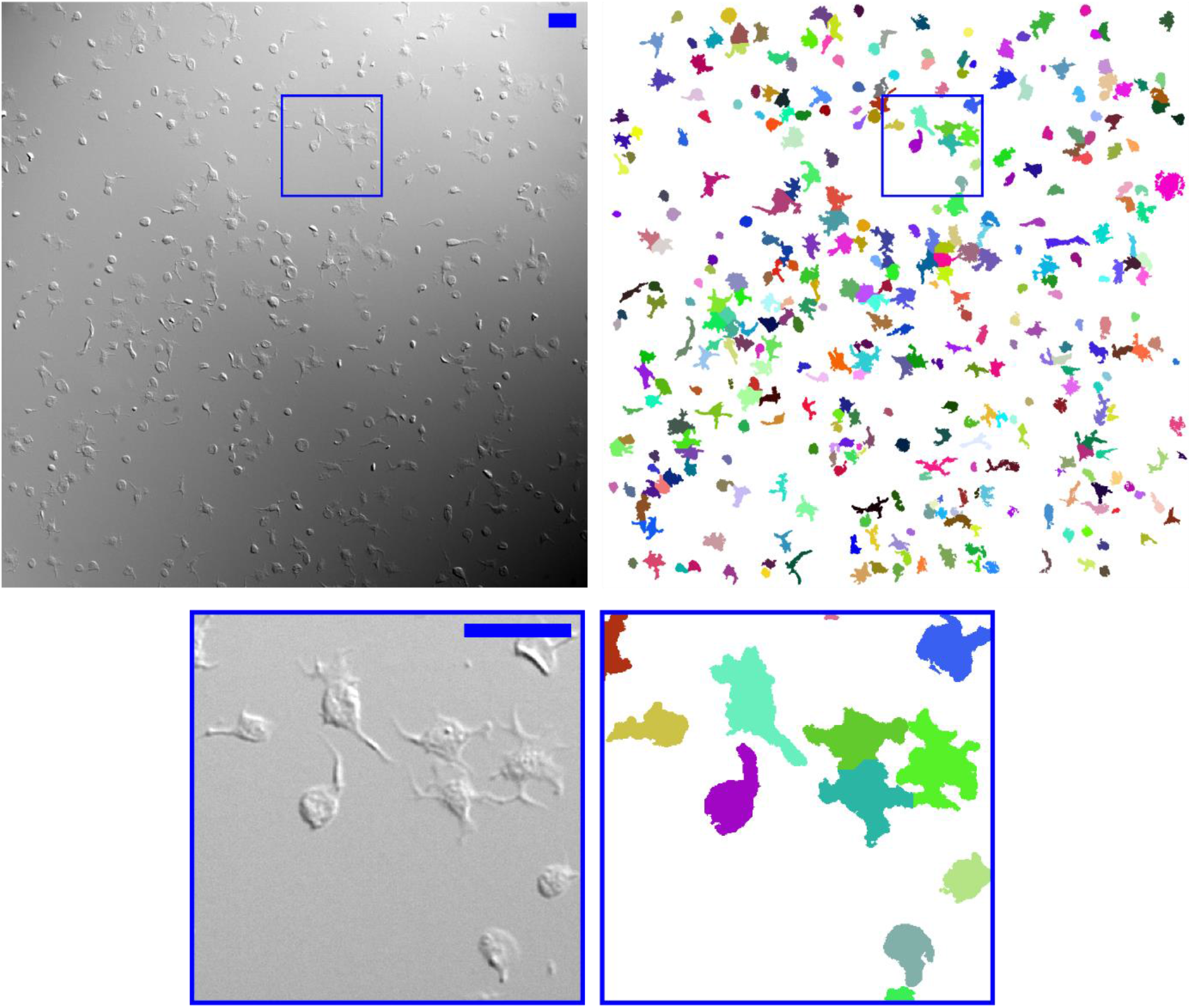
Phase contrast images and segmentation results for a representative acquisition where platelets were spread on fibrinogen. Full field of view and cropped images are shown. Each cell segmentation is represented by a different colour. The workflow is able to produce reasonable segmentation without using a florescent stain. Scale bar 10 μm.

